# CNNcon: A Quantitative Imaging Tool for Lung CT Image Feature Analysis

**DOI:** 10.1101/615492

**Authors:** Jason Causey, Jake Qualls, Jason H. Moore, Fred Prior, Xiuzhen Huang

## Abstract

**Background:** Lung CT scans are widely used for lung cancer screening and diagnosis. Current research focuses on quantitative analytics (radiomics) to improve screening and detection accuracy. However there are very limited numbers of portable software tools for automatic lung CT image analysis.

**Results:** Here we build a Docker container, CNNcon, as a quantitative imaging tool for analyzing lung CT image features. CNNcon is developed from our recently published algorithm for nodule analysis, based on convolutional neural networks (CNN). When provided with a list of the centroid coordinates of regions of interest (ROI) in a volumetric CT study containing potential lung nodules, CNNcon can automatically generate highly accurate malignancy prediction of each ROI. CNNcon can also generate a vector of image features of each ROI, to facilitate further analyses by combining image features and other clinical features. As a Docker container, CNNcon is portable to various computer systems, convenient to install, and easy to use. CNNcon was tested on different computer systems and generated identical results.

**Conclusions:** We anticipate that CNNcon will be a useful tool and broadly acceptable to the research community interested in quantitative image analysis.

**Availability:** CNNcon and document are publicly available and can be downloaded from the website: http://bioinformatics.astate.edu/CNN-Container/

## 1 Background

Lung cancer is the leading cause of cancer related death for both men and women worldwide [http://www.who.int/news-room/fact-sheets/detail/cancer]. In the United States, lung cancer was diagnosed in 222,500 people in 2017 and accounted for an estimated 155,870 deaths [Siegel2017]. Early detection and diagnosis of lung cancer is critical to give patients the best chance at recovery and survival. Lung CT scans are broadly used for lung cancer screening and diagnosis.

Significant recent research effort has focused on combining quantitative image analysis and machine learning (radiomics) to improve the speed and reliability of lung cancer screening and early detection [Kalpathy-Cramer2016a; 2016b; Lambin2012]. Much of this work has been conducted under the auspices of NCI’s quantitative imaging network (QIN), which emphasizes the creation and validation of reliable quantitative imaging tools [Clarke2014].

In an assessment of the state of imaging informatics in precision medicine published by members of the QIN, the need for reliable, validated radiomics analysis tools deployed using container technology was emphasized [Chennubhotla2017]. However currently there are very few such portable software tools available to the community. Especially there are few Docker containers for automatic lung CT image feature analysis. We conducted Pubmed and Google Scholar searches with related key words, such as, “docker and image feature”, “docker and lung CT”, “docker and CT”, “CNN docker and CT”. We found that the various searches returned only a few relevant publications; For example, for the search with key words “docker and CT”, only one relevant publication [Echegaray2017] was found. The Quantitative Image Feature Engine (QIFE) is a framework for creating radiomics pipelines which has been deployed in containerized form. Thus far this tool has been used to generate and analyze engineering features not features learned from the data using deep learning techniques. We found one other publication from a research group in Spain, which was focused on a problem of CT reconstruction, and the work compared two approaches for managing the software dependencies of the code: store the software libraries on a Storage Element and using containers for executing the job [Chillarón2017].

We have developed a Docker container, CNNcon, as a quantitative imaging tool for automatically analyzing lung CT image features utilizing our recently published algorithms [Causey2018]. When provided with a list of the centroid coordinates of regions of interest (ROIs) in a volumetric CT study containing potential lung nodules, CNNcon can automatically generate a malignancy prediction of each ROI, as well as generate a vector of image features of each ROI. As a Docker container, CNNcon is can be easily deployed and is easy to use.

## 2 Implementation

An automation script was created to automate the steps necessary to isolate the region of interest (ROI) corresponding to each centroid in the input centroids list. Running the automation script automatically performs the following steps:

- A temporary storage directory is created at the specified location if it does not already exist.
- For each centroid in the input centroids list, the corresponding DICOM image is loaded, then a region surrounding the centroid is cropped from the original image and stored along with necessary metadata in an intermediate file in the HDF5 format.
- The HDF5 intermediate file containing all ROIs is loaded, and each ROI is evaluated by the selected NoduleX [Causey2018] CNN model.
- An output file containing, for each ROI, either probabilities corresponding to the likelihood of the ROI belonging to the “positive” class, or feature vectors representing the ROI according to the CNN model’s feature space are output in a CSV-compatible format.

A “Dockerfile” was created to provide instructions for Docker [https://www.docker.com] to package the necessary code components and operating system environment into a suitable Docker container. This file was used to produce a Docker image that will be referred to as nodulex henceforth. This Docker image may be run from the command line as shown in the procedures.

The container is designed to be run as a command-line evaluation tool, or as a step in a larger analysis pipeline. The environment must be prepared before running the tools as follows:

- Input DICOM files should be placed into a directory (here referred to as “data”). Each individual scan must be in a subdirectory of the data directory, with the name of the subdirectory corresponding to a unique identifier for the scan, i.e. the patient ID or scan ID.
- A list of the centroid coordinates corresponding to regions of interest (ROIs) must be prepared in a comma-separated (CSV-compatible) format. The centroid list must be stored in a directory that will be available to the running container, such as in the data directory. Here we refer to this file as centroid_list.csv and assume it is stored in data, i.e. its path is data/centroid_lists.csv.
- A directory must be designated to receive the results output by the analysis. Here we refer to this directory as results.
- A temporary directory to support the necessary intermediate files must be specified. (Intermediate files require approximately 70KiB per ROI with default settings.) Here we refer to this directory as tmp.

The directory structure as described above would look like the following:

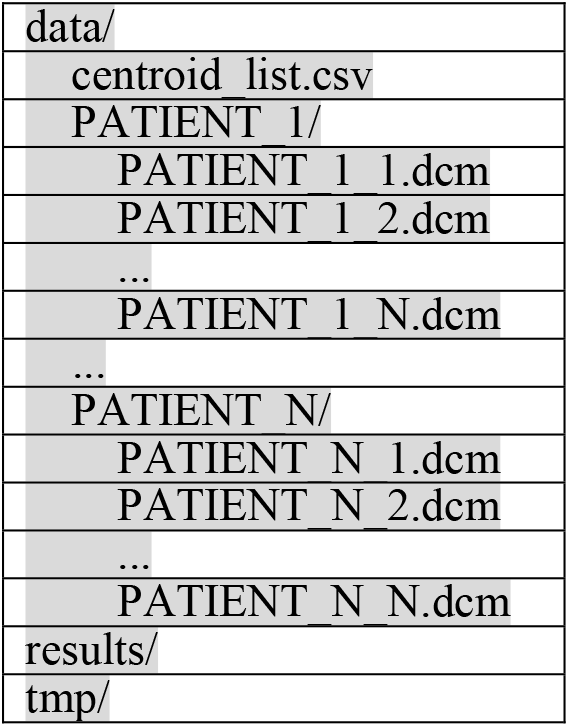

|  |
| --- |
| data/ |
| centroid_list.csv |
| PATIENT 1/ |
| PATIENT 1 1.dcm |
| PATIENT 1 2.dcm |
| ... |
| PATIENT 1 N.dcm |
| ... |
| PATIENT N/ |
| PATIENT N 1.dcm |
| PATIENT N 2.dcm |
| ... |
| PATIENT N N.dcm |
| results/ |
| tmp/ |

The centroid file is used to describe both IDs and locations for each of the regions of interest. The file must be formatted as follows:

- The first line of the file may be a header line; if so, it must begin with the # character.
- Each line of the file corresponds to one ROI

– Fields are separated by commas; no other commas may appear in the data.
– The Patient ID (or Scan ID) is the first field on the line; it must correspond exactly to the name of the subdirectory in data that contains the DICOM files for that scan.
– The second field is the slice number (or Z-axis coordinate) of the center of the ROI.
– The third field is the X-axis coordinate (or row number) of the center of the ROI.
– The fourth field is the Y-axis coordinate (or column number) of the center of the ROI.
– There may be a fifth field that is a label for the ROI class; if so, it will be ignored by this procedure.

An example of the centroids file format is shown below:

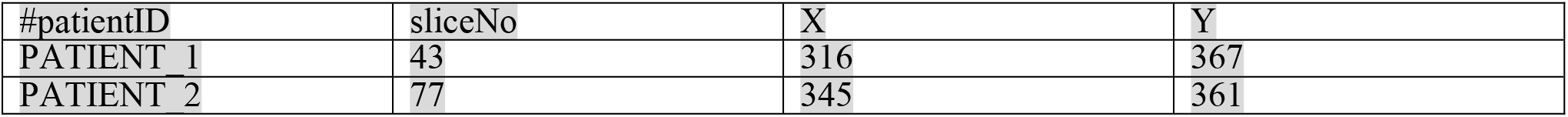

| #patientID | sliceNo | X | Y |
| --- | --- | --- | --- |
| PATIENT 1 | 43 | 316 | 367 |
| PATIENT 2 | 77 | 345 | 361 |

### Procedure 1: Predictions with default parameters

Assuming the directory structure is as shown above, the following Docker command will execute the analysis procedure with default parameters and produce the result file predictions.csv in the results directory. The results file will contain predictions in the range [0,1] corresponding to the probability that the ROI is in the “positive” class. With default parameters, the “positive” class corresponds to the LIDC-IDRI “malignancy” classification ≥4[armato2011lung].

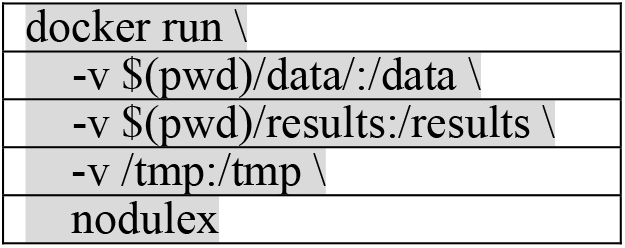

Note that in this example $(pwd) is used to make the path to the data and results directories absolute, as is required by Docker, and the default system temporary directory /tmp is used for tmp. It is necessary to modify the paths to suit the host system’s configuration. Additionally, on some systems it is necessary to have superuser permissions to run Docker, requiring the use of sudo (replace docker with sudo docker in the command shown).

### Procedure 2: Feature vectors with default parameters

Using the same assumptions as in Procedure 1, this command will produce the result file expression.csv in the results directory. The results file will contain a 200-dimensional vector for each ROI corresponding to the “feature layer” in the model’s neural network (the last fully-connected layer prior to the softmax classification layer).

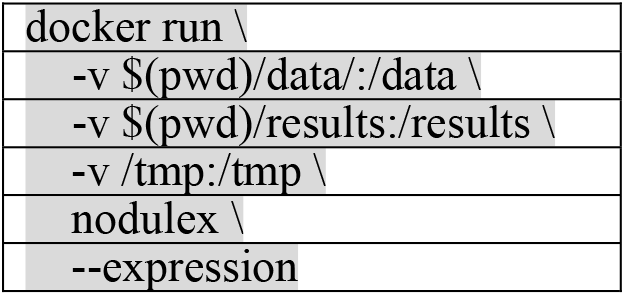

### Listing of command parameters

The container supports several optional parameters, as explained below.

### Optional positional argument

centroid-file:

File defining the centroid coordinates of ROIs to examine (CSV-like format). Must be in a directory accessible to the container (i.e. data, tmp, or results).

### Optional arguments

-h, --help:

Displays information about available options and exits.

-c, --cache:

Flag indicating the tools should cache intermediate results in the temporary directory, leaving them intact for future use.

-d DATA, --data DATA:

Directory (relative to the container) containing the DICOM data, with one subdirectory per patient, named according to patient ID.

-r RESULTS, --results RESULTS:

Directory (relative to the container) into which results will be written.

-t TMP, --tmp TMP:

Directory (relative to the container) where intermediate and cache data will be written.

-s SIZE, --size SIZE:

Voxel size (in X,Y dimensions) of the input region. Supported values are 21 and 47. The default is 47.

-x, --expression:

Store “expression” feature vectors in output directory instead of “prediction” probabilities. The output file will be “expression.csv”.

-m MODEL, --model MODEL:

Selects which model will be used for classification. Supported values are “12v45”, “1v45”, “NvNN”. The default is “12v45”.

-v VOXEL_SHAPE, --voxel-shape VOXEL_SHAPE:

Specifies voxel shape in mm. A scalar value indicates homogeneous voxels (X=Y=Z), or use a list of three numbers i.e. “X,Y,Z” to specify non-cubic voxels. Use the flag “−1” to keep original shape from input DICOM image unchanged. The default is “−1”.

-w, --window:

Flag indicating the model should use window normalization during preprocessing. This is the default when –size is 47.

-n, --no-window:

Flag indicating the model should not use window normalization during preprocessing (the model will use min-max normalization instead). This is the default when –size is 21.

## 3 Results

The Docker container CNNcon was tested on two different computers with two different settings: the default setting and the setting with the option of a specified homogeneous voxel. For each of the two settings, based on our testing with 132 modules, the two computers with different operating systems and hardware platforms generated identical results. *Please refer to http://bioinformatics.astate.edu/CNN-Container/ for the files, including the input csv file, the output prediction file and the feature vector file*. The following is the details of the testing.

We analyzed 132 nodules from the LIDC-IDRI study, consisting of 66 in the “positive” class with malignancy ratings 4 or 5, and 66 in the “negative” class with malignancy ratings 1 or 2. We recorded accuracy and area under the Receiver Operating Characteristic curve (AUC) for the predictions. With default settings, the model produced predictions with an accuracy of 82.6%, corresponding to an AUC of 0.896. The 132 nodules from 121 CT images were processed in 46 minutes, 45 seconds on a Dell OptiPlex 5050 with a 3.6GHz Intel Core i7 CPU and 16GB of RAM running Ubuntu Linux. The same dataset was processed in 71 minutes, 9 seconds on a 2016 MacBook Pro with 2.9 GHz Intel Core i7 CPU and 16GB of RAM, with identical results.

Additionally, we analyzed the same 132 nodules again, demonstrating the option to specify a homogeneous voxel shape. We chose the voxel shape (x,y,z) = (0.7,0.7,1.0) mm. With this setting, the model’s accuracy was increased to 84.1% corresponding to an AUC of 0.914. This experiment ran in 57 minutes, 57 seconds on the Dell Optiplex 5050 and in 82 minutes, 57 seconds on the 2016 MacBook Pro with identical results. The increase in processing time is due resampling the images to the common voxel shape.

## 4 Discussion

We developed the Docker container CNNcon, as a quantitative imaging tool for automatically analyzing lung CT image features. CNNcon can automatically generate malignancy prediction of each ROI, and also generate a vector of image features of each ROI. CNNcon was tested on two different computers and generated identical results; and when tested with two different settings, the generated results are also consistent. Advantages of using Docker containerization for the analysis pipeline include ease of installation, portability, and simplified analysis procedure. We plan to deploy CNNcon net on two different high performance computing platforms and on a commercial cloud to further characterize its portability and performance. We anticipate that CNNcon, as a Docker container, will be a useful tool and broadly acceptable to the research community.

### Funding

This work was partially supported by NIH NCI U01CA187013, and NSF 1452211, 1553680, and 1723529, NIH R01LM012601, as well as partially supported by NIH P20GM103429.

### Contributions

X.H., J.C., and F.P. conceived and designed the study. J.C., J.Q. J.M., F.P. and X.H. implemented the design, analyzed the data, evaluated the software through numerical testing, and worked on manuscript preparation. X.H. and J.C. wrote the paper, with input from all authors.

